# Olfactory perception of chemically diverse molecules

**DOI:** 10.1101/049999

**Authors:** Andreas Keller, Leslie B. Vosshall

**Author notes:** (AK), (LBV).

## Abstract

Background: Understanding the relationship between a stimulus and how it is perceived reveals fundamental principles about the mechanisms of sensory perception. While this stimulus-percept problem is mostly understood for color vision and tone perception, it is not currently possible to predict how a given molecule smells. While there has been some progress in predicting the pleasantness and intensity of an odor, perceptual data for a larger number of diverse molecules are needed to improve current predictions. Towards this goal, we tested the olfactory perception of 480 structurally and perceptually diverse molecules at two concentrations using a panel of 55 healthy human subjects.

Results: For each stimulus, we collected data on perceived intensity, pleasantness, and familiarity. In addition, subjects were asked to apply 20 semantic odor quality descriptors to these stimuli, and were offered the option to describe the smell in their own words. Using this dataset, we replicated several previous correlations between molecular features of the stimulus and olfactory perception. The number of sulfur atoms in a molecule was highly correlated with the descriptors “garlic” “fish” “decayed,” and large and structurally complex molecules were perceived to be more pleasant. We discovered a number of strong correlations in intensity perception between molecules, which suggests a shared mechanism for perceiving these molecules. We show that familiarity had a strong effect on the ability of subjects to describe a smell. Many subjects used commercial products to describe familiar odors, highlighting the role of prior experience in biasing verbal report of perceived smells. Nonspecific descriptors like “chemical” were applied frequently to unfamiliar smells, and unfamiliar odors were generally rated as neither pleasant nor unpleasant.

Conclusions: We present a very large psychophysical dataset and use this to correlate molecular features of a stimulus to olfactory percept. Our work reveals robust correlations between molecular features and perceptual qualities, and highlights the dominant role of familiarity and experience in assigning verbal descriptors to smells.

## INTRODUCTION

In olfaction, the conscious percept of a smell is often discussed in terms of perceived intensity, perceived pleasantness, and perceived olfactory quality (“garlicky” “flowery” etc.). The perceived intensity of a stimulus is the most basic and least ambiguous of these measures. Previous research has shown that only sufficiently volatile and lipophilic molecules are odorous [1]. Molecular features such as molecular weight or the partial charge on the most negative atom correlate with perceived intensity. Several of these molecular features were used in regression equations that modeled the intensity of 58 molecules with impressive accuracy (R^2^=0.77-0.79) [2]. However, to our knowledge, this prediction has not been tested in an independent dataset and a formal model that relates chemical structure to intensity has yet to be developed [3]. Several models have been developed to predict perceived pleasantness of an odorant based on its physical features [4-6]. Both molecular size [4, 6] and molecular complexity [5] correlate with perceived pleasantness. There are also well-known predictions of olfactory quality. Molecules containing sulfur atoms are predicted to smell “garlicky,” whereas molecules containing ester groups are predicted to have a “fruity” smell. However, to our knowledge, these predictions of individual olfactory qualities have not yet been quantified.

Two features of olfactory perception complicate solving the stimulus-percept problem in this sensory modality. The first complication is that different individuals perceive the same molecules with different sets of functional odorant receptors [7-11]. These differences have been shown to influence perception [8, 11-15], and the same molecule is therefore often perceived differently by different individuals. This complication is not unique to olfaction. Colorblind individuals perceive the same visual stimulus differently from standard observers. However, in olfaction, the variability between different individuals is unusually large [16-18]. The second feature of olfaction that has to be considered when attempting to predict perception based on stimulus features is that prior experience, cultural practices, motivational state, and non-olfactory information affect verbal reports of olfactory perception. The common co-occurrence of sweet tastes and odors that are described as smelling “sweet,” for example, has led to the suggestion that odors such as vanillin acquire their sweet smell quality by being experienced together with sweet tastes [19].

Furthermore, olfactory psychophysics suffers from a paucity of empirical data necessary to formulate theories to relate stimuli to percept. Many past attempts to solve the stimulus-percept problem for olfaction have used the same dataset from Andrew Dravnieks, who carried out a study in which expert panelists evaluated 138 different molecules using 146 standard semantic descriptors [20]. The purpose of the Dravnieks study was to develop a standard lexicon for describing olfactory stimuli of interest to the flavor and fragrance industry. Accordingly, both the molecules themselves and the semantic descriptors attached to them represent only a small number of possible odors and percepts that humans can experience. Although there are alternative sources of data on the perceptual qualities of larger numbers of molecules, these are often based on the judgments of experts from companies that provide fragrance materials [21, 22]. Information from these sources is not standardized, and it can be difficult to assess how the data were obtained and how reliable they are. These constraints have slowed attempts to relate the molecular structure of an odorant to its conscious percept by human subjects.

To improve current predictions, perceptual data for a larger number of diverse molecules is needed. In this study, we present and analyze data on the perception of 480 structurally diverse molecules at two concentrations. The molecules evaluated by our subjects are more diverse than those used in previous studies. Another improvement of our dataset is that we provide individual responses in addition to the average perception of the group of subjects, allowing us to avoid masking perceptual variability. The motivation behind producing this dataset was to increase the number and diversity of molecules that can be used to test formal models that predict perceived smell based on features of the molecules. All raw data are being made freely available with the publication of this work to stimulate further analysis by other groups.

We found that intensity perception was strongly related to vapor pressure and molecular weight. We also uncovered strong correlations in intensity perception between certain pairs or clusters of variably intensity-rated stimuli, suggesting that these odorants may be sensed by the same odorant receptors. The presence of sulfur atoms biased molecules to be perceived as unpleasant. Conversely, pleasantness was correlated with molecular complexity. Finally, we discovered that familiarity strongly biases olfactory perception. Unfamiliar stimuli were less likely to receive a semantic descriptor and tended to be neither pleasant nor unpleasant. This suggests that semantic categorization of olfactory stimuli alone is unlikely to solve the stimulus-percept problem.

## RESULTS

We tested the perception of 480 different molecules at two concentrations in 61 healthy subjects. 20 molecules were tested twice at both concentrations, for a total of 1,000 stimuli tested. The molecules ranged in molecular weight from 18.02 (water) to 402.54 (tributyl-2-acetylcitrate) with a median of 144.24 (Figure 1a), and in molecular complexity from 0 (water and iodine) to 514 (androstadienone) with a median of 109 (Figure 1b). Molecular complexity is a rough estimate of a molecule’s complexity that considers the variety of elements and structural features of the molecule [23]. Many of the molecules had unfamiliar smells. Of the stimuli that subjects could perceive, 70% were rated as unknown and were given low familiarity ratings (Figure 1c, left), while those rated as known had high familiarity ratings (Figure 1c, right). The molecules were structurally and chemically diverse (Figure 1d), and some have never been used in prior psychophysical experiments. The 480 molecules had between 1 and 28 non-hydrogen atoms, and included 29 amines and 45 carboxylic acids. Two molecules contained halogen atoms, 53 had sulfur atoms, 73 had nitrogen atoms, and 420 had oxygen atoms.

**Figure 1.**
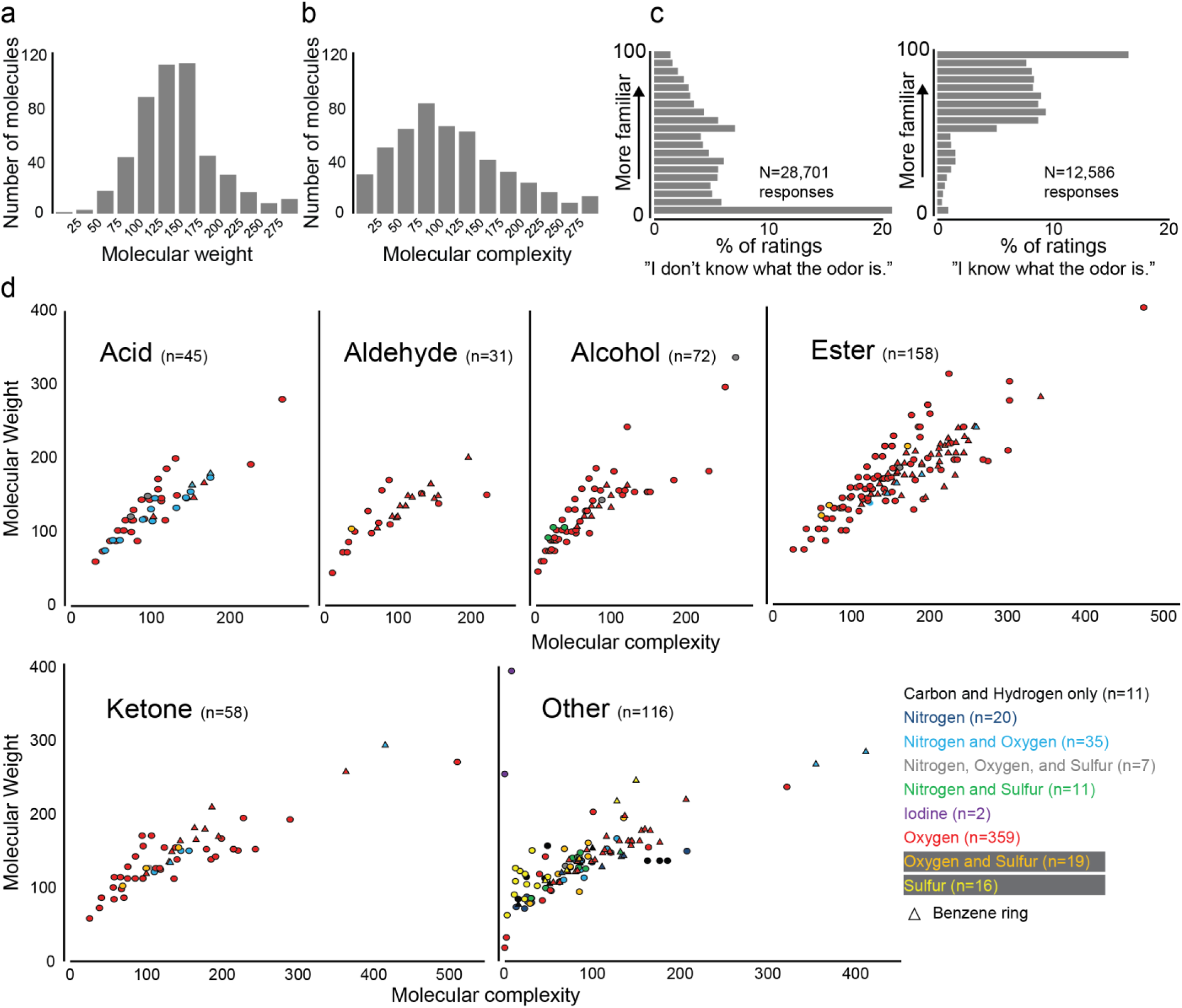
Molecules. **a-b**, Molecular weight (a) and molecular complexity (b) of the molecules used in this study. **c**, Histograms of familiarity ratings (0-100, binned in 20 units of 5) for stimuli that subjects identified as unknown (left) or known (right). N denotes the total number of responses across all stimuli and all subjects, **d**, Molecular weight and molecular complexity parsed by chemical functionality.

The 1,000 stimuli were tested across 10 visits, and the order of visits and stimuli within visits were randomized. The sequence of prompts for each stimulus is shown in Figure 2a. For questions about familiarity, intensity, pleasantness, and the 20 descriptors, subjects were presented with a slider that they moved along a line. The final position of the slider was translated into a scale from 0 to 100. The 20 descriptors were “edible” “bakery” “sweet” “fruit” “fish” “garlic” “spices” “cold” “sour” “burnt” “acid” “warm” “musky” “sweaty” “ammonia/urinous” “decayed” “wood” “grass” “flower” “chemical”.

**Figure 2.**
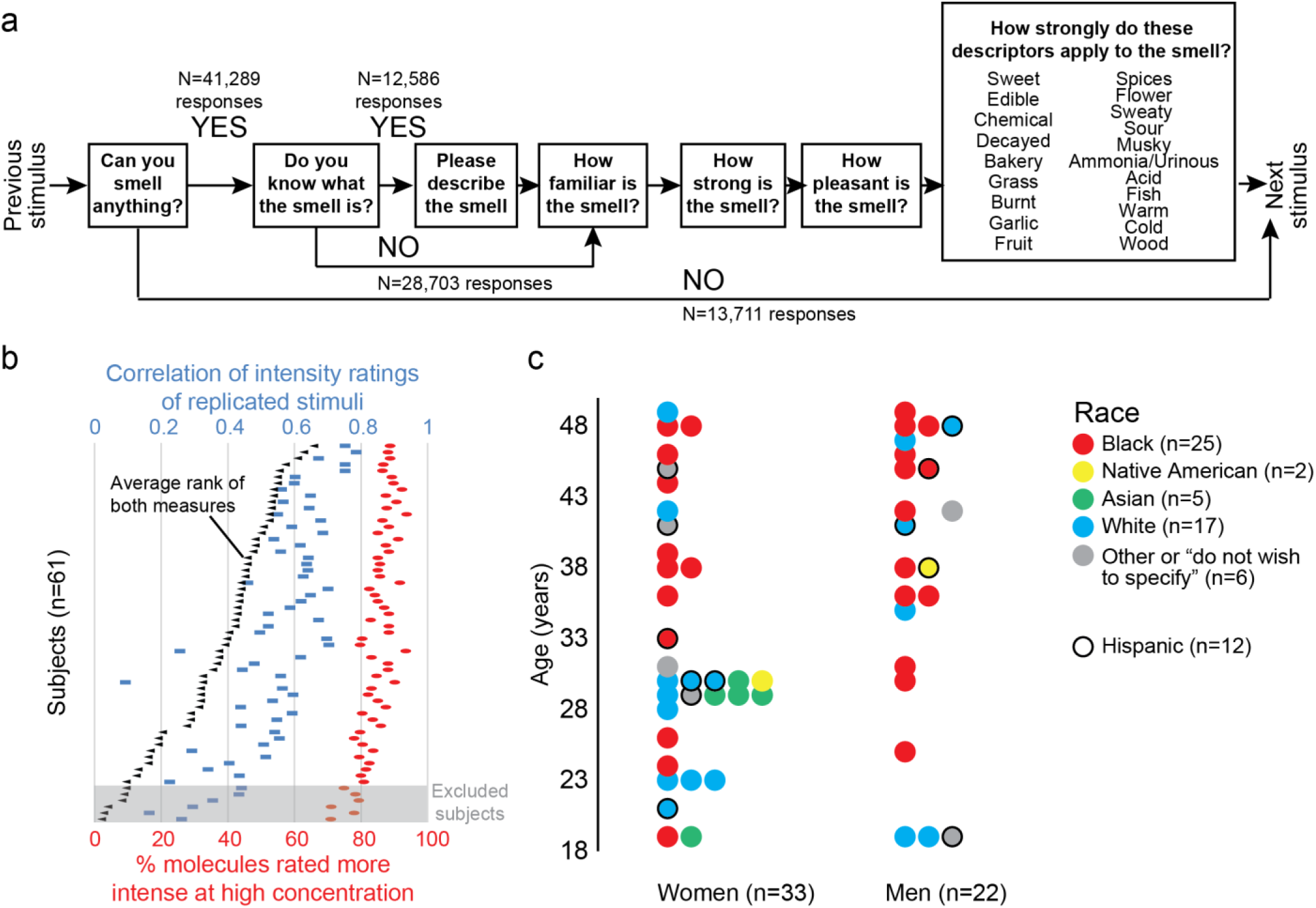
Subjects. **a**, Sequence of prompts for each stimulus. N denotes the total number of responses across all stimuli and all subjects, **b**, General olfactory performance of the 61 subjects who completed the study. Six subjects with the lowest rank in replicability of intensity ratings were excluded from further analysis, **c**, Age, gender, and self-reported race and ethnicity of 55 evaluated subjects.

61 subjects completed all ten visits. To exclude malingerers and subjects suffering from hyposmia, the average rank of two objective measures of olfactory performance was calculated and the 6 lowest ranked subjects were excluded from further analysis (Figure 2b), and data from the remaining 55 subjects formed the basis of all analysis in the paper (Figure 2c) (Additional file 1). Twenty arbitrarily chosen molecules were presented twice at both concentrations throughout the study. In general, intensity and pleasantness (Figure 3a) and descriptor usage (Figure 3b) were consistent between the two presentations. For reasons that we do not understand, the intensity ratings and descriptors for high concentrations of 2-methyl-1-butanol (odor 2), isopropyl acetate (odor 13), and thiophene (odor 19) differed substantially between the two presentations (Figure 3a-b).

**Figure 3.**
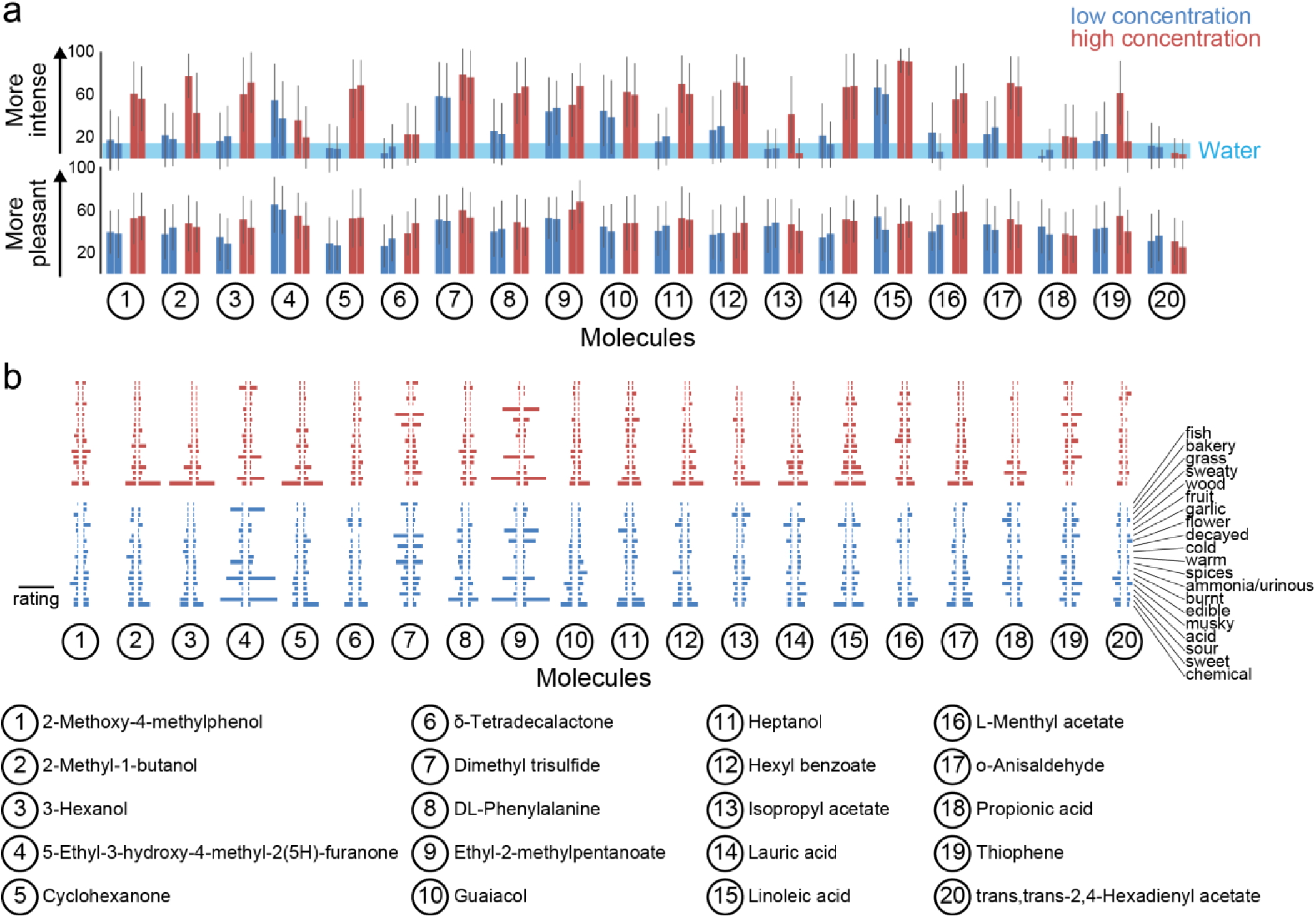
Repeated stimuli. **a**, Ratings for intensity (top) and pleasantness (bottom) for the 40 stimuli (20 molecules at two concentrations) each presented twice (mean ± S.D.). **b**, Ratings of descriptors for high (top) and low (bottom) concentrations of 20 molecules each presented twice. Average ratings of descriptors for first (left-facing bar plot) and second (right-facing bar plot) presentations. Scale bar: rating of 50 on a scale of 0 to 100.

### Perception of the stimuli

437 of the 1,000 stimuli were presented at the same dilution (1/1,000). Among these, methyl thiobutyrate was rated to be most intense, followed by 2-methoxy-3-methylpyrazine, 2,5-dihydroxy-1,4-dithiane, butyric acid, and diethyl disulfide (Figure 4a). On the opposite end of the intensity scale, 61 molecules were rated to be less intense than the average intensity rating of the two dilutions of water (14.44) when diluted 1/1,000 (Additional file 1). In the majority of cases, stimuli presented at high concentration received higher intensity ratings than the same stimuli at low concentration (Figure 4b).

**Figure 4.**
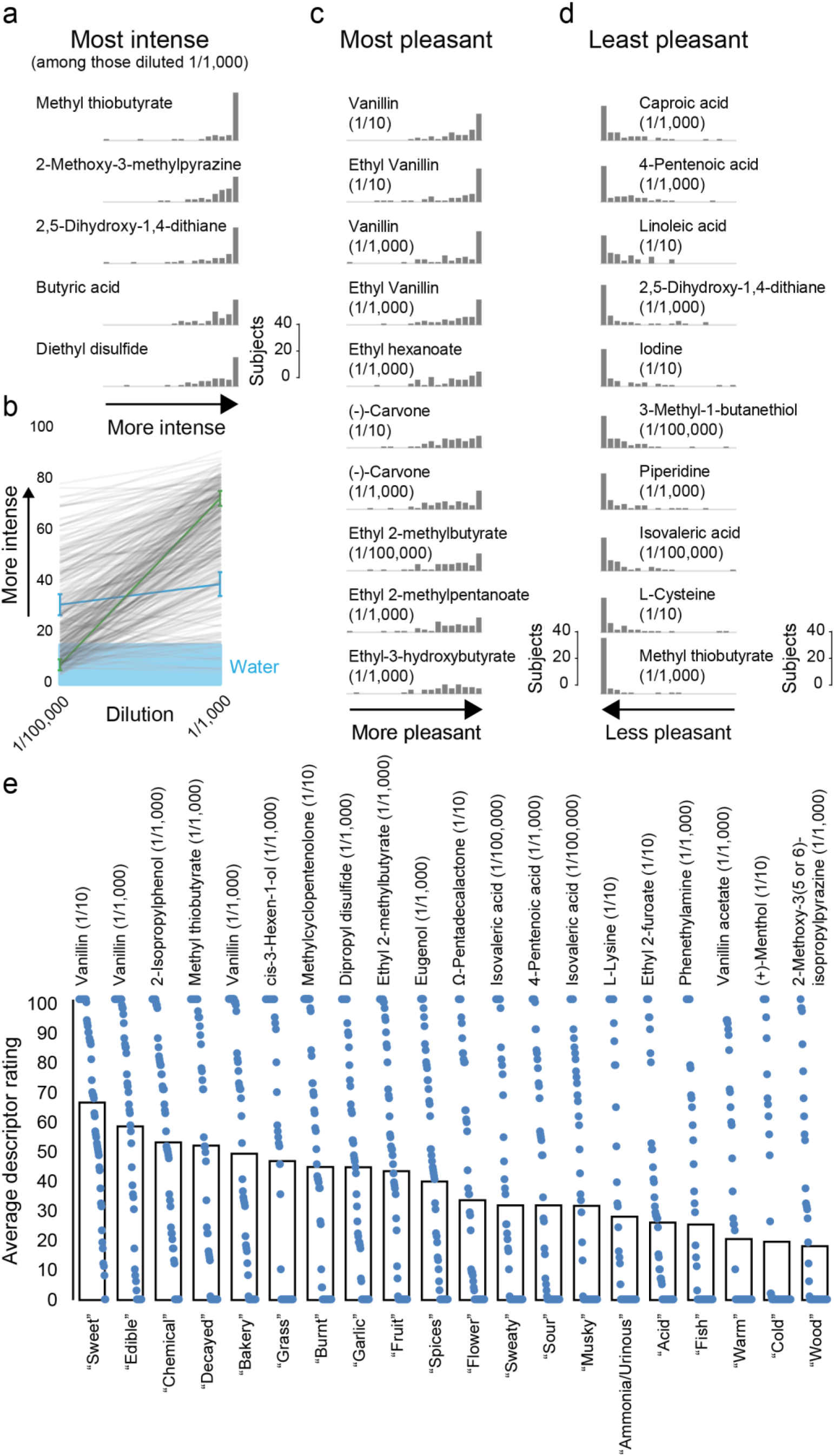
Perception of stimuli. **a**, Histograms of intensity ratings for the 5 most intense of 437 stimuli presented at 1/1,000 dilution (most intense on top). **b**, Average intensity ratings of 437 molecules presented at 1/1,000 and 1/100,000 dilutions, with standard error of the mean shown for two molecules [methyl salicylate (green) and methyl caprylate (blue)]. **c-d**, The 10 most pleasant (most pleasant on top) (**c**) and 10 least pleasant of the 1,000 stimuli (least pleasant on bottom)(**d**). **e**, Descriptor rating of the stimuli most representative of each of the 20 descriptors. Open bars show average ratings, and blue dots indicate individual ratings. Only the 778 stimuli perceived to be more intense than water (14.44) were included in this analysis. In **a**, **c-d**, histograms of subject ratings of intensity (**a**) or pleasantness (**c-d**) are plotted on a scale from 0-100, binned in 20 units of 5.

In addition to perceived intensity, we tested perceived pleasantness. The two concentrations of vanillin and ethyl vanillin accounted for the four most pleasant stimuli in the study. Both concentrations of (-)-carvone and four different esters comprised the remainder of the ten most pleasant stimuli (Figure 4c). The least pleasant stimulus was methyl thiobutyrate, which was also the most intense of the stimuli diluted 1/1,000 (Figure 4a). Another three of the ten least pleasant stimuli were also sulfur-containing molecules and four others were carboxylic acids (Figure 4d).

The frequency by which a descriptor is attached to a molecule reveals molecules that are representative of each of the descriptors (Figure 4e). Methyl thiobutyrate (1/1,000), the most intense and least pleasant stimulus, received the most ratings of “decayed”. Vanillin received the highest rating for “edible” (1/1,000), “bakery” (1/1,000), and “sweet” (1/10), and vanillin acetate (1/1,000) was rated the “warmest” stimulus. Isovaleric acid (1/100,000) received the highest rating for both “musky” and “sweaty” (Figure 4e).

### Variability in intensity perception

While some stimuli were nearly unanimously rated as very intense (Figure 4a), and many others were perceived to be very weak by all subjects, a few stimuli showed great variability in intensity ratings (Figure 5a). Androstadienone (1/1,000), which is known to be perceived differently by different subjects [8, 14, 17], was the stimulus with the third most variable intensity perception in this dataset behind benzenethiol and 3-pentanone (Figure 5a). In some cases, there was a correlation between the intensity perception of different molecules, which may point towards a shared mechanism for perceiving these molecules. The four stimuli pairs with the strongest correlation between the perceived intensity ratings are shown in Figure 5b. Strong correlations such as these can be caused by a ceiling effect in which two odors that are perceived to be very intense by all subjects will show a strong correlation that may not be indicative of a shared perceptual mechanism. However, the correlation between 5-methylfurfural and propyl acetate is an example of a strong correlation between two stimuli whose intensity perception varies between subjects.

**Figure 5.**
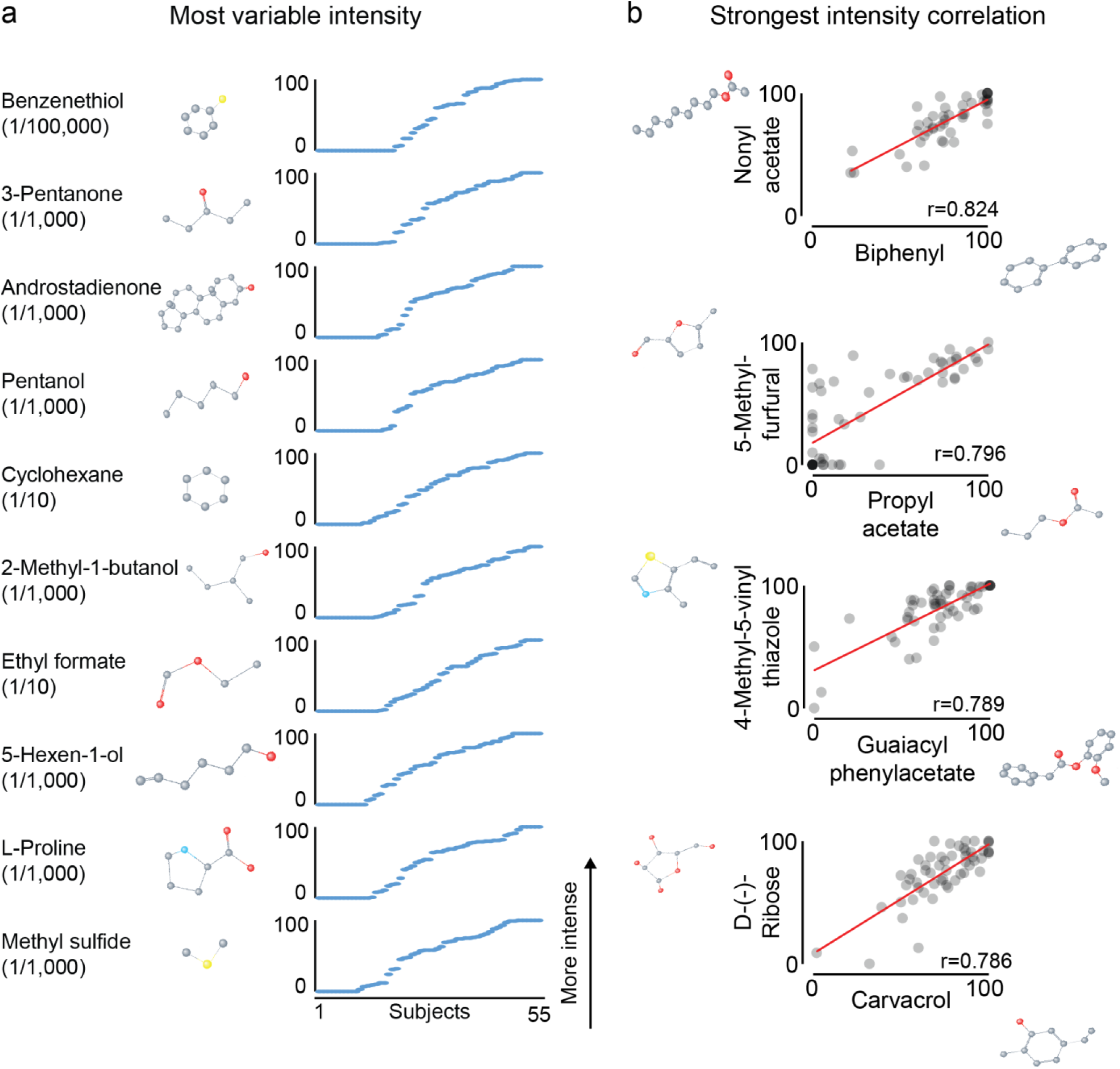
Variability in perception. **a**, The 10 stimuli with the most variability in intensity (most variable on top). **b**, The 4 pairs of all stimuli with the largest correlation between intensity ratings. Only the 778 stimuli perceived on average to be more intense than water (14.44) were included in this analysis.

### Descriptor usage and familiarity

In addition to providing intensity and pleasantness ratings, subjects rated whether and how strongly each of 20 semantic descriptors applied to the stimuli. The most commonly used descriptor was “chemical” and the least frequently used descriptor was “fish” (Figure 6a). The strategies by which different subjects applied descriptors to stimuli varied considerably. One subject used only 508 descriptors for the 1,000 stimuli, whereas another used 9,678 descriptors (Figure 6b, left). This is consistent with previous reports that descriptor usage frequency is an individual trait [20]. The median number of total descriptors applied was 1,900, meaning that the average subject applied around two descriptors to the average stimulus. Different subjects also applied individual descriptors with different frequency (Figure 6b, right). Three subjects did not apply the descriptor “fish” to any of the stimuli. At the other extreme, one subject applied this descriptor to more than half of the stimuli (505/1000). At the median, subjects applied “fish” to 21 of the stimuli. The two outliers who applied “fish” to more than 150 stimuli were also the two subjects who applied the largest number of descriptors overall. The frequency of applying the descriptor “chemical” also varied between subjects. One subject applied it to only 80 stimuli, another to 798 stimuli.

**Figure 6.**
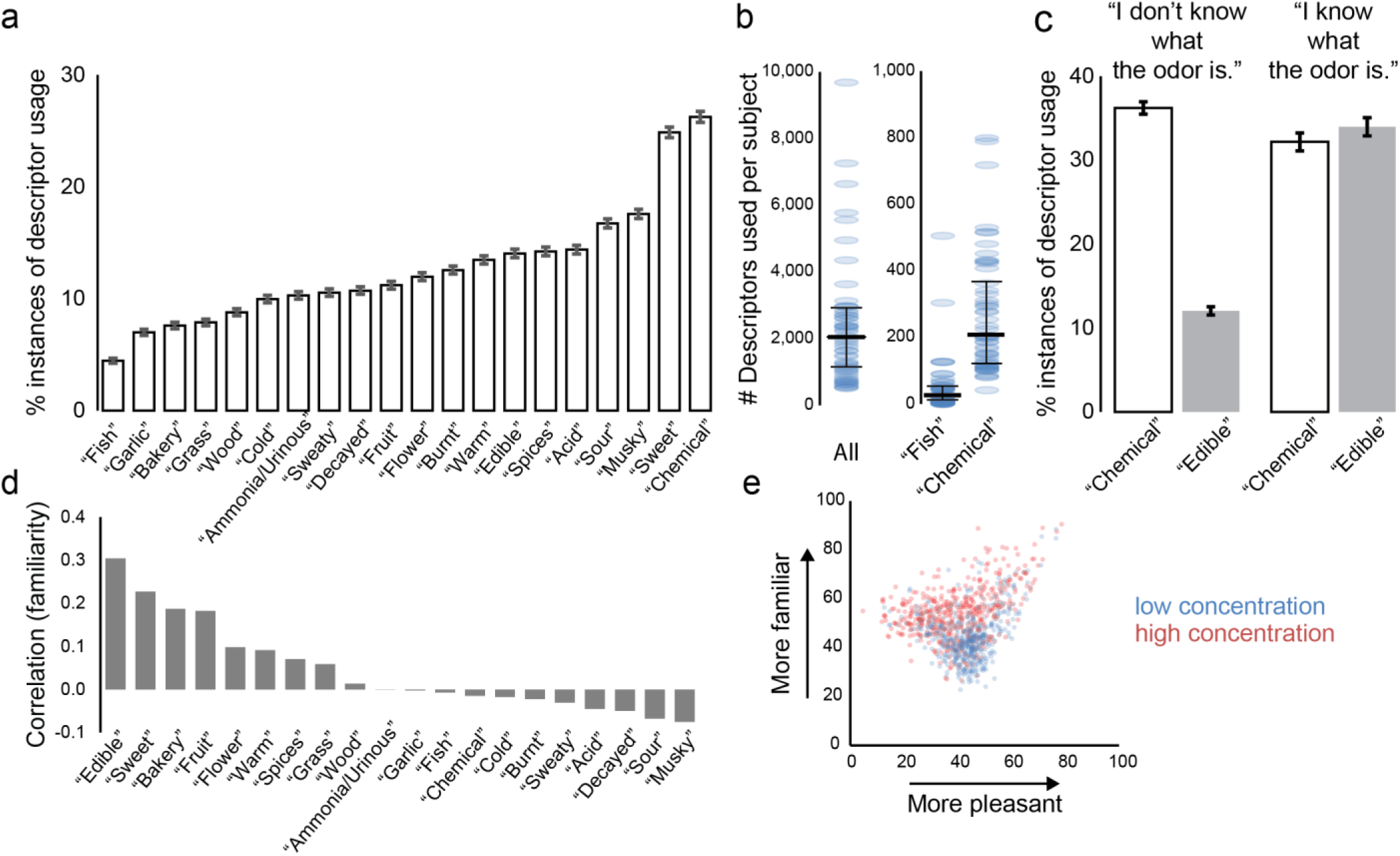
Descriptor usage and familiarity. **a**, Descriptor usage for all subjects (with 99% confidence interval indicated). 100% would correspond to a descriptor assigned to all stimuli by all subjects. **b**, Descriptor usage per subject. Left: all descriptors (maximum possible value: 20,000). Right: “chemical” “fish” descriptors (maximum possible value: 1,000). Data from 55 individual subjects (blue) and median and first and third quartiles (black). **c**, Descriptor usage for “chemical” and “edible” for all stimuli (99% confidence interval indicated), with responses divided according to unknown (left: N=28,703 responses) and known (right: N=12,586 responses) stimuli. 100% would correspond to a descriptor assigned to all stimuli by all subjects. **d**, Correlation between familiarity ratings and the ratings of 20 descriptors. **e**, Average familiarity and pleasantness ratings for 1,000 stimuli.

Some descriptors were predominantly applied to stimuli the subjects were familiar with, whereas others were often used for unfamiliar smells. For example, for unfamiliar smells “chemical” was a more common descriptor than “edible” (Figure 6c, left), whereas both were used equally for familiar stimuli (Figure 6c, right). Correlations between familiarity and the ratings for the 20 descriptors showed that “edible” was most strongly correlated with high familiarity (Figure 6d). Perceived pleasantness also had an interesting relationship with familiarity (Figure 6e). Unfamiliar stimuli tended to be neither pleasant nor unpleasant, whereas the most pleasant stimuli were also judged to be very familiar.

### Correlations between descriptors

The descriptors used in this study do not refer to independent qualities of smells. Stimuli that were perceived as “fruity” were more likely to be perceived as “sweet,” perhaps because many fruits are sweet. As can be seen in Figure 7a, the descriptors “sweet” “flower” “edible” “fruit” “bakery” were strongly correlated with a high rating for pleasantness. In contrast, the descriptors “decayed” “musky” “sour” “sweaty” were correlated with a low pleasantness rating. Between the descriptors, the highest correlation was between “edible” and “bakery”, followed by “sweet” and “fruit”, “sweet” and “edible”, and “musky” and “sweaty”. On the other hand, some descriptors were mutually exclusive and therefore negatively correlated. The strongest negative correlation was found between “edible” and “chemical”, followed by “sweet” and “musky”, and “sweet” and “sweaty”.

**Figure 7.**
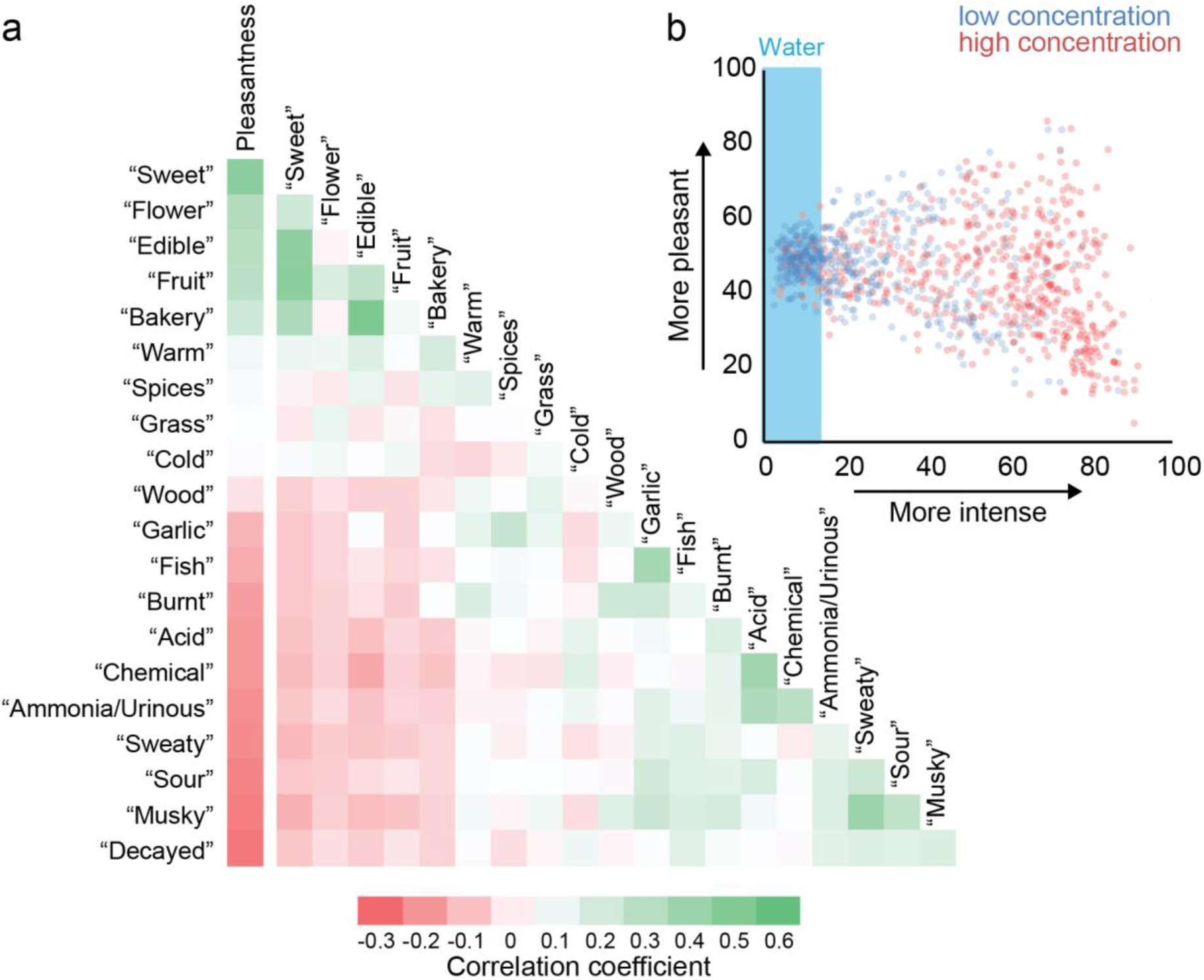
Correlations between descriptors. **a**, Heat map of correlation between pleasantness ratings and the ratings of 20 descriptors. **b**, Average intensity and pleasantness ratings for 1,000 stimuli.

None of these negative correlations between descriptor pairs was as strong as the correlation between pleasantness and intensity (Figure 7b). Very unpleasant stimuli tended to be perceived as very intense. However, as can be seen in Figure 7b, the relationship between perceived intensity and pleasantness is more complex than this. Weak stimuli were perceived as neither very pleasant nor very unpleasant. Both the 29 most unpleasant and the 9 most pleasant stimuli had an intensity rating over 50.

### Subjects’ own words

In addition to rating intensity, pleasantness, and 20 descriptors, subjects were given the opportunity to describe the stimuli in their own words (Figure 2b and Additional file 1). Overall, the words used by the subjects were subsets of descriptors used by fragrance professionals [20]. Words such as “sweet” “burnt” “grass” “candy” “vanilla” were common (Figure 8a). However, there were also some idiosyncrasies. Women tended to describe more of the stimuli than men (Figure 8b). Providing self-generated descriptors was optional, and subjects used their own words to describe between 2 and 803 of the 1,000 stimuli (median = 173). The subject who provided descriptors for 2 stimuli used only 2 total words (2 unique words: “burnt,” “paint”). The subject who described the most stimuli used 7160 words, 766 of them unique. The most common words used by this subject were “sweet,” “pencil,” and “eraser,” which were used 156, 139, and 133 times, respectively. Both the subject that described the least and the subject that described the most stimuli had above average performance as determined by the metrics in Figure 2b, suggesting that the variability in the number of described stimuli is due to behavioral rather than perceptual variability. Subject-generated descriptions were similar to published descriptors found in the Sigma-Aldrich Flavor and Fragrance catalogue, on Wikipedia, or in Dravnieks’s odor atlas (Figure 8c). One notable difference was that subjects used product names such as Vicks VapoRub^®^, Marshmallow Fluff^®^, and Bengay^®^, to describe smells (Figure 8c). As expected, few subjects attempted to describe the smell of water (Figure 8c).

**Figure 8.**
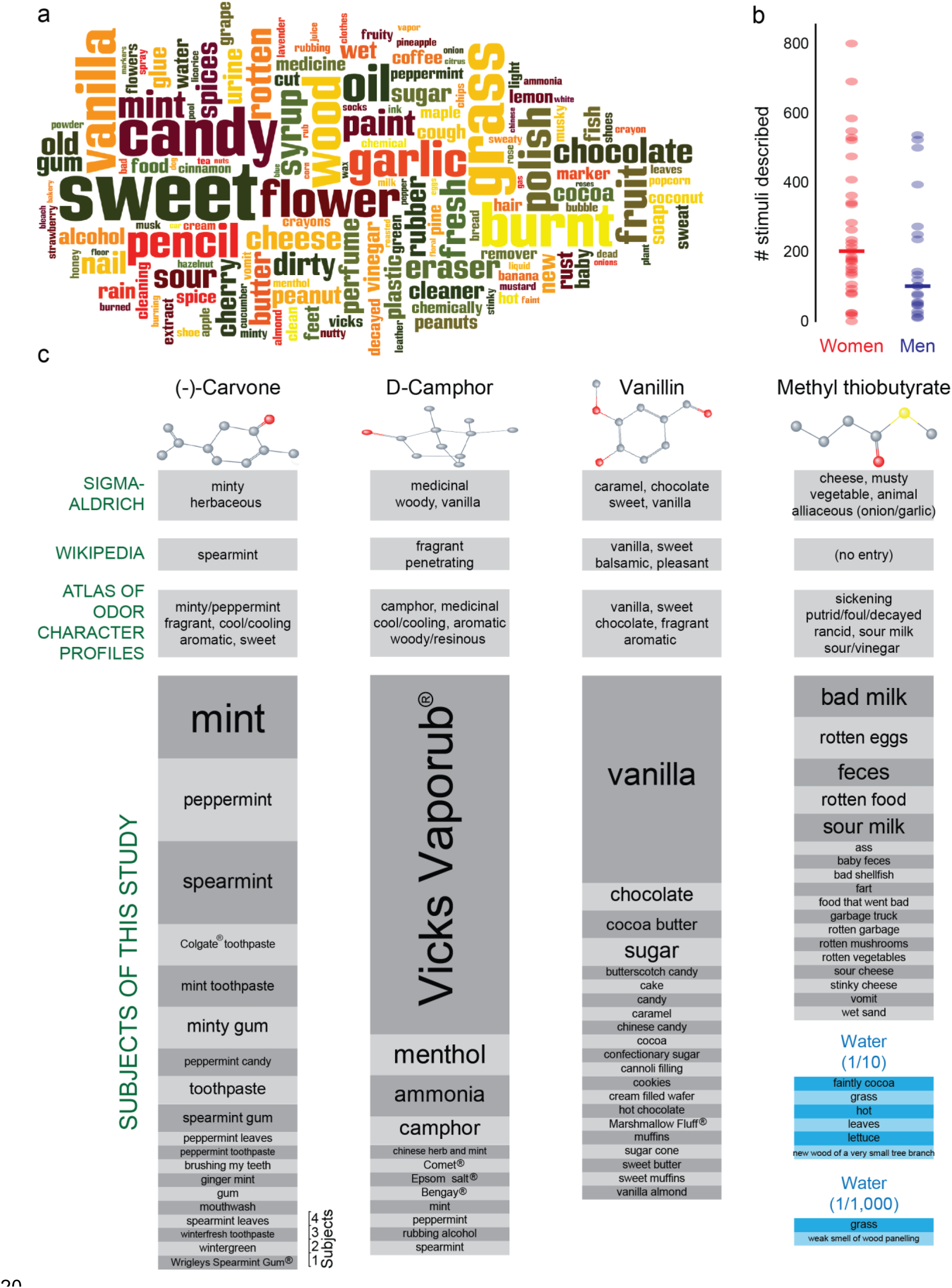
Subjects’ own words. **a**, A word cloud in which font size represents the frequency with which words describing odor quality were used. **b**, The number of stimuli that each of the 55 subject described in their own words. Individual data are shown as dots, median as line. c, Semantic odor descriptors for (-)-carvone (1/10), D-camphor (1/10), vanillin (1/10), and methyl thiobutyrate (1/1,000). Published descriptors from Sigma-Aldrich Flavor and Fragrance Catalogue, Wikipedia, and the five descriptors with the highest applicability from the Dravnieks odor atlas [20] (top) and self-generated descriptors provided by subjects for the same 4 odor stimuli as well as water “diluted” 1/10 or 1/1,000 (bottom).

### Structure-odor relationships

The dataset presented here makes it possible to investigate the relationship between the physical features of molecules and their perceptual qualities. This is illustrated in Figure 9a, which shows the physical features of the molecules that have the strongest positive correlation with the ratings of intensity, pleasantness, and each of the 20 descriptors. The most basic perceptual quality in any modality is the intensity with which the stimulus is perceived. 437 stimuli in this study were diluted 1/1,000, and this subset of stimuli can be used to investigate which physical features predict the perceived intensity of a molecule. We found a positive correlation between the vapor pressure of a molecule and its intensity (Figure 9b, top). The molecular feature that had the strongest positive correlation with perceived intensity of the stimuli diluted 1/1,000 in this study is the presence of an atom-centered fragment that contains a sulfur atom (Figure 9a). We also replicated the previously reported negative correlation between molecular weight and perceived intensity (Figure 9b, bottom) [2].

**Figure 9.**
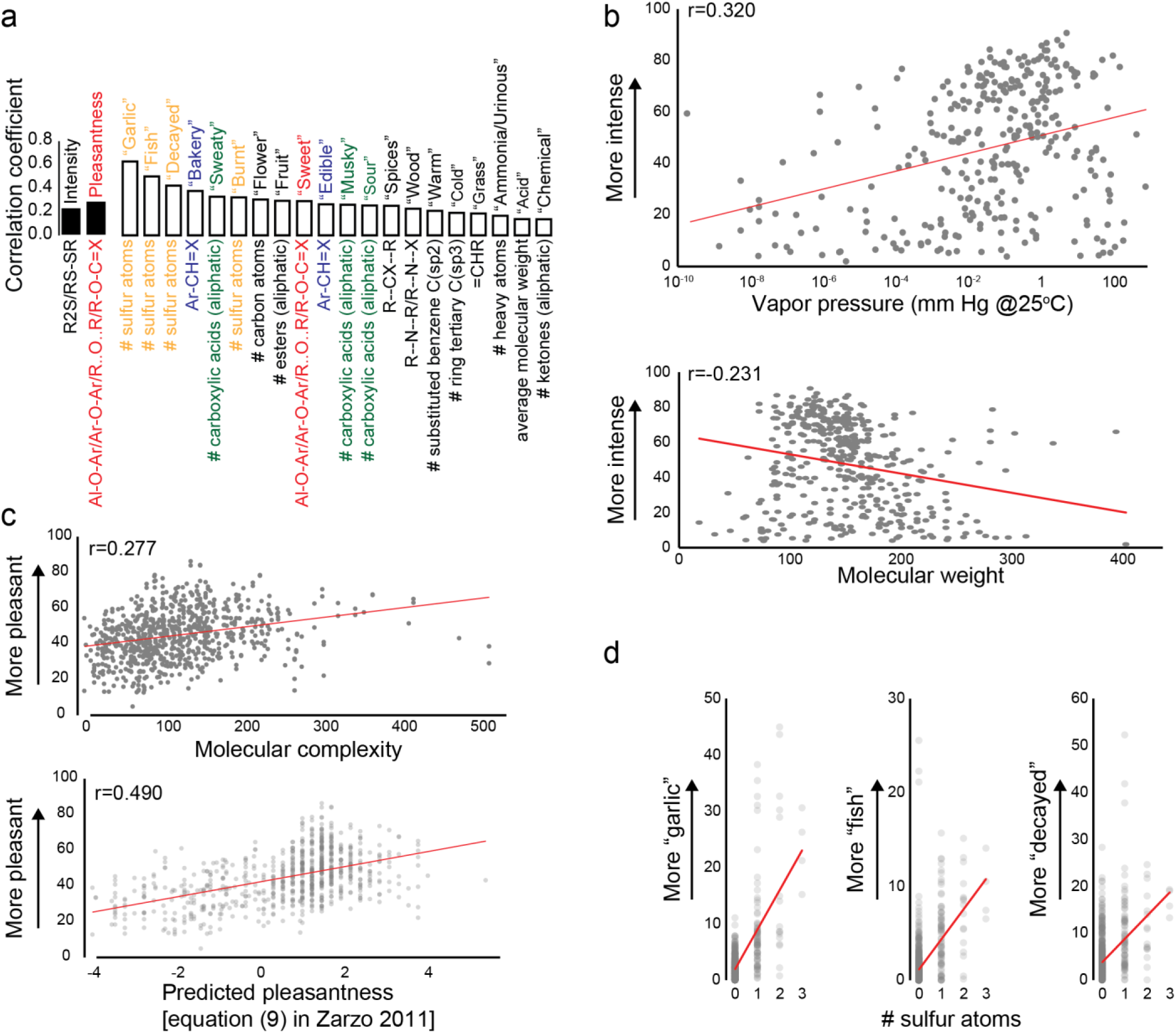
Predicting perception. **a**, The strongest positive correlations between a molecular feature and intensity, pleasantness, and descriptor ratings. **b**, Perceived intensity and vapor pressure (top; limited to the 319 molecules with available vapor pressure information) and perceived intensity and molecular weight (bottom). **c**, Pleasantness and molecular complexity (top), and pleasantness and molecular features from equation (9) in [4]: −2.62+0.23*number of atoms (excluding H)+1.58*presence of oxygen-1.96*presence of sulfur-2.58*presence of an acid group-1.89*presence of an amine group (bottom). **d**, The number of sulfur atoms and ratings for “garlic” (left), “fish” (middle), and “decayed” (right). In all panels, only stimuli diluted at 1/1,000 are included in analysis of intensity so that only stimuli diluted to the same level are compared; and only the 778 stimuli perceived to be more intense than water (14.44) were included in the analysis of pleasantness.

Another perceptual quality of smells that has been predicted using molecular features is pleasantness [6]. The proposal that molecules with higher molecular complexity are more pleasant [5], was replicated by our dataset (Figure 9c, top). Another prediction of perceived pleasantness was based on the observation that the number of atoms (excluding H) and the presence of an oxygen atom result in molecules that are perceived to be more pleasant, whereas the presence of sulfur, an acid group, or an amine group make it less pleasant [4]. This prediction was also replicated by our dataset (Figure 9c, bottom).

The goal of most research that attempts to link perceptual and molecular determinants of a smell is to predict whether certain semantic descriptors are likely to be applied to the smell of a molecule. Our data show that the extent to which a given descriptor correlates with molecular features differs between descriptors (Figure 9a). For the descriptors “chemical” and “acid” there were only weak correlations with chemical features. The strongest correlations were seen between the descriptors “garlic” “fish” “decayed” and the number of sulfur atoms in the molecule (Figure 9d).

## DISCUSSION

We present here a dataset that captures the sensory perception of 480 different molecules at two different concentrations as experienced by diverse population of human subjects. Subjects rated intensity, pleasantness, familiarity, and applied 20 odor descriptors. 40 stimuli (20 molecules at two concentrations) were presented to the subjects twice, and there was generally robust test-retest reliability. Subjects were capable of matching intensity ratings to the concentration of the stimulus molecules. 98% of the molecules that subjects perceived to be stronger than water (intensity rating of 14.44) were perceived to be more intense at the high concentration than at the low concentration (median intensity rating difference: 33.2).

We discovered a strong influence of familiarity on the semantic description of odors. In addition, the descriptors applied to the stimuli replicated how descriptors were applied by experts in many cases [20]. Among the molecules used both in this study and in Dravnieks’s study, diethyl disulfide was the most representative of the descriptor “garlic” in both studies. Similarly, the molecules most representative of “flower” (2-phenylethanol), “decayed” (methyl thiobutyrate), “sweaty” (isovaleric acid), and “spicy” (eugenol), were the same in the two studies. For some other descriptors like “sweet” and “sour,” there was also large overlap in descriptor usage for given molecules.

However, we also found marked differences in how descriptors were used by our untrained subjects and experts. For example, subjects used “musky” to describe unpleasant body odors. In contrast, experts generally use “musky” to describe compounds naturally sourced from animal glands or their synthetic analogues. These are often used as base notes in perfumery, and experts associate musks with pleasant descriptors such as “animalic” “sweet” “powdery” “creamy.” However for our subjects, “musky” had a strong negative correlation with pleasantness, and was strongly correlated with the descriptor “sweaty”. The molecule rated as most “musky” in this study was isovaleric acid, which experts do not rate as “musky” [20]. The five molecules that Dravnieks lists as representative of the “musk” descriptor are also rated “fragrant” and “perfumery” by experts [20]. Therefore, the word “musky” has a colloquial meaning that is different from its technical meaning in perfumery.

### A new dataset for human psychophysics research

This new dataset differs from most other sources of information about how different molecules are perceived. First, our dataset includes molecules that are usually considered to be odorless, like water, glycerol, and citric acid. We included these stimuli to create data on an outgroup that so far has been missing from olfactory research. In addition to including odorless molecules in our dataset, we included molecules with unfamiliar smells not easily associated with descriptors commonly used to classify smells. As a consequence, subjects did not recognize more than 69% of the stimuli that they could perceive. In contrast, many other datasets consist largely of molecules that are representative of specific descriptors. This practice leads to the danger that language conventions used to describe odors are studied instead of odor perception itself. Avery Gilbert and Mark Greenberg succinctly summarized the dangers of this approach when they wrote that “we are creating a science of olfaction based on cinnamon and coffee” [24] (page 329). Including odorless molecules and smells that do not align well with common semantic descriptors allows for a more comprehensive analysis of human smell perception.

A second, related, feature of our dataset is that it includes a rating of the familiarity of the stimulus. In datasets like those by Dravnieks [20] or Arctander [22], in which molecules were largely chosen because of their economic and cultural importance, the experts that evaluated the odors very likely knew what they were smelling. We believe that it is important to study the perception of unfamiliar smells alongside familiar smells. The danger of studying smells that are easily identified is that the responses of the subjects conflate qualities of the smell with qualities of what the smell is associated with. If a smell is identified as banana smell, the pleasantness rating will not only reflect how pleasant the smell is, but also whether the subject likes eating bananas or not. In contrast, smells that elicit no associations are evaluated based exclusively on perceptual qualities. Familiarity ratings are a way to determine whether the ratings reflect perceptual qualities, or whether they reflect a mix of perceptual qualities and associations with those qualities.

A third feature that distinguishes this dataset from many other sources of information about olfactory perceptual qualities is that we not only report a population average, but also how individual subjects rated the stimuli. This information is useful because olfactory perception depends not only on the molecular features of the stimulus, but also on the perceptual system of the perceiving subject. Olfactory perceptual systems differ considerably between individuals [7-11].

To stimulate further analysis of the data in this study, we are making the entire dataset freely available (Additional file 1). Our analysis here is primarily concerned with predicting how different molecules are perceived. However, the dataset enables the investigation of other topics, for example differences in perception between different demographic groups [18]. Perceptual correlations between stimuli (Figure 5b) can be used to arrange molecules in a perceptual odor space, or to investigate the underlying mechanisms of shared perception.

### Predicting intensity

There are two complications to predicting perceived intensity. First, the intensity of a given molecule at a given dilution is not only dependent on the interaction between the molecule and the perceptual system, but also on how many of the molecules will reach the odorant receptors. How many molecules will be released from the dilution depends on the molecule’s vapor pressure and solubility in the solvent. The vapor pressures in Figure 9b are for the pure molecule and do not take into account interactions with the solvent. Once the molecules arrive at the olfactory epithelium, the efficiency with which they reach the hydrophobic binding pockets of odorant receptors depends on their lipophilicity. Consequently, perceived intensity has a high positive correlation with vapor pressure (r=0.320) and a high negative correlation with a measure of hydrophilicity (squared Moriguchi octanol-water partition coefficient (logP^2^); r=-0.321).

The second complication is that different molecules have different stimulus response functions, which determine the relationship between the dilution of the molecule and its perceived intensity [25]. Because the relation between dilution and perceived intensity differs between different molecules, it is possible that a molecule perceived to be stronger than another molecule at one dilution will be perceived as weaker than the other molecule at a different dilution. This is clearly illustrated by the two molecules highlighted in Figure 4b. While methyl salicylate was perceived to be more intense than methyl caprylate at the 1/1,000 dilution, methyl caprylate was perceived to be more intense than methyl salicylate at the 1/100,000 dilution. Because of this complication, it is not enough to predict the perceived intensity of a molecule at a given dilution. Instead, the parameters of the stimulus response function have to be predicted. The two dilutions used for the molecules in this study are not sufficient to determine the parameters of the stimulus response functions, but they allow for more sophisticated prediction of perceived intensity than those based only on a single dilution or on the molecule’s detection threshold.

The 437 molecules that were presented in this study at a dilution of 1/1,000 were perceived to be of very different intensities. 63 of the 437 molecules were perceived to have lower intensity than water (intensity rating 14.44), but at the other extreme the 1/1,000 dilution of methyl thiobutyrate and other molecules were perceived to be very strong stimuli. The previously reported correlation between vapor pressure and perceived intensity as well as the negative correlation between molecular weight and perceived intensity were reproduced in this dataset [2]. The data are clearly more complex than correlations between single molecular features and perceived intensity can capture. For example, low perceived intensity was reported with molecules of very low and very high vapor pressure. We also discovered that a single molecular feature, the presence of a certain sulfur-containing atom-centered fragment, has a positive correlation to perceived intensity that is almost as strong as the correlation between molecular weight and intensity.

### Predicting pleasantness

In this work we replicated the finding that perceived pleasantness correlates with molecular complexity [5]. We also confirm the observations that perceived pleasantness correlates with the number of atoms (excluding H) and that the presence of an oxygen atom results in molecules that are perceived to be more pleasant, whereas the presence of sulfur, an acid group, or an amine group make it less pleasant [4]. A third prediction for pleasantness [6] that made it possible to predict pleasantness with r~0.5 was partially based on molecular features that were not provided by the version of the cheminformatic software we used (Dragon 6) [26, 27]. However, the chemical features from the model in [6] that our version of the software also provided, like the number of non-H atoms, showed the same relation with pleasantness in our data set as in [6].

The molecular feature that we discovered here to have the strongest correlation with pleasantness is the presence of a certain atom-centered fragment containing an oxygen atom. Our data show that three previous models that predict pleasantness performed well on our independent dataset, and suggest that a combination of these models might outperform any single model.

### Predicting odor qualities

Most research into structure-odor relationships is concerned with explaining why a given semantic descriptor, such as “musky” or “camphorous” is commonly applied to some molecules, but not to others [1, 28, 29]. Among the 20 descriptors used here, the strongest correlation between a descriptor and a molecular feature was between the semantic descriptor “garlic” and the number of sulfur atoms in the molecule. The correlation coefficient for this correlation is 0.63. In contrast, the strongest positive correlation of the descriptor “chemical” with any molecular feature is 0.14. The differences in the strength of correlations between semantic descriptors and molecular features suggest that the application of some semantic descriptors (“garlic” “fish”) can easily be predicted based on molecular features, whereas the application of others (“chemical” “acid”) either is much more complex or cannot be predicted based on molecular features.

A plausible explanation for this observation is that all semantic descriptors that are assigned to smells must be learned by association. It may be that the situations in which subjects formed an association between the word “garlic” and a specific smell are very similar between subjects. They may have been formed when the subject was eating a meal with a lot of garlic in it that was described to them as smelling like “garlic”. The associations between the word “chemical” and a specific smell on the other hand are probably different between subjects. All molecules are chemicals and the descriptor “chemical” is used by subjects to describe a wide variety of molecules. In our study, this descriptor was the default in cases in which the subject could not identify the smell. The perceptual diversity of molecules that are described as “chemical” is illustrated by the fact that Dravnieks lists as the five molecules that are representative of the descriptor “chemical” phenyl acetylene (sweet, floral), anisole (anise seed-like), pyridine (fishy), cyclohexanol (camphor-like), and 1-butanol (banana-like, alcoholic) [20].

What this example suggests is that some descriptors are specific, whereas others are ambiguous and open to different interpretations. In other words, some descriptors have a single reference smell, with which they are strongly associated whereas other descriptors have several or no reference smells. This results in weak associations between the descriptor and the reference smells that vary between individuals.

#### Conclusions

Psychophysical data with a larger number of chemically diverse molecules will increase our understanding of the relationship between stimulus and perception in olfaction. The dataset presented here reproduces findings from previous studies, confirming that it is a useful dataset for such a project. However, it is overly simplistic to think that the perception of molecules could be predicted entirely based on physical features of the molecules. How a molecule is perceived is determined by both the perceptual system and the physical features of the stimuli. In vision, the same light stimulus is perceived differently by monochromatic, dichromatic, and trichromatic subjects. In olfaction, the receptors that detect odorants vary greatly between individuals [7-11], and this variability leads to differences in how the same molecule is perceived by different subjects [8, 11-15]. The dataset presented here makes it possible to make predictions for individual subjects because we not only report average or consensus perception, but also how each individual subject perceived the stimuli. The data presented here also reveal that the assignment of descriptors depends strongly on familiarity with the smell. We presume that subjects only applied a given descriptor to a stimulus when they could retrieve the memory of the reference smell that they associated with the descriptor. The specificity of the reference smell depends on the descriptors. “Spices” is a semantic descriptor that can trigger a variety of different smell associations, whereas “garlic” refers to a more specific type of smell. We anticipate that only descriptors with an unambiguous reference odor can be predicted based on molecular features.

Another problem with verbal descriptors is that they are culturally biased. The current standard set of 146 Dravnieks descriptors was developed in the United States in the mid-1980s and is increasingly semantically and culturally obsolete. Even if these descriptors were updated to be current and relevant across different nationalities and cultures, it is unlikely that semantic descriptors will ever cover to totality of olfactory perceptual space. Moreover, because the existing descriptors were developed with a small list of stimuli, new untested molecules or complex mixtures of molecules may lack appropriate semantic descriptors. To circumvent the limitations of verbal descriptors, an alternative semantic-free approach to predict similarity between stimuli based on molecular features [30] should be pursued. Initial implementations of this method have shown astonishing success, producing a correlation of r=0.85 between predicted and empirically-determined stimulus similarity [30]. Predicting perceptual similarity between olfactory stimuli would result in a complete and comprehensive ability to predict the perception of any molecule. The usage of verbal descriptors can then be predicted by predicting the similarity of a stimulus to a signature odor that is representative of that descriptor.

Studies that aim to predict smell perception based on molecular features have been given a boost by the introduction of Dragon software, which calculates thousands of different molecular features [26, 27]. This large collection of molecular features frees researchers from guessing what features of molecules influence how they are perceived, and makes it possible to test a wide variety of molecular features to find those that play a role in determining a molecule’s smell. However, the large number of molecular features available for building formal models of structure-odor relationships also brings the danger of overfitting. An overfitted model fits the data that was used to create it well, but it has poor predictive performance. Overfitting often occurs when a model has too many parameters relative to the number of observations. Reducing the likelihood of overfitting by increasing the number of molecules that can be used to test formal models was a major motivation behind generating this dataset.

Importantly, we have used a subset of the data presented here for a competitive modelling challenge in collaboration with IBM Research, Sage Bionetworks, and DREAM challenges [31]. For this competition, predictive models were built based on a training set containing one set of stimuli and then evaluated using a test set containing a different set of stimuli. The DREAM Olfaction Prediction Challenge aims to develop the most comprehensive computational approach to date to predict olfactory perception molecule smells based on the physical features of a molecule. The results of this challenge will soon be published elsewhere.

## METHODS

### Ethics, consent, and permissions

All behavioral testing with human subjects was approved and monitored by The Rockefeller University Institutional Review Board (protocol LVO-0780). Subjects gave their written informed consent to participate in these experiments.

### Subjects

Healthy subjects between the ages of 18 and 50 were recruited from the New York City metropolitan area and tested between February 2013 and July 2014. 61 subjects completed all 10 study visits. The remaining subjects dropped out before all 10 visits were completed, or were not invited back after the first visit at our discretion. To exclude malingerers and hyposmic subjects, the average rank of two objective measures of olfactory performance was calculated and the 6 lowest ranked subjects were excluded from further analysis (Figure 2b). The measures were the correlation between intensity rankings of 40 repeated stimuli (Figure 3a, top and Figure 2b, blue symbols), and the number of molecules that subjects rated as more intense at the high versus low concentration (Figure 4b and Figure 2b, red symbols). The 6 subjects with the lowest average rank of both measures were excluded (Figure 2b, black symbols), leaving 55 subjects (33 female) whose data comprise the results of this paper. Of these, 25 self-identified as black, 17 as white, 5 as Asian, and 2 as Native American. 12 self-identified as Hispanic (Additional file 1). The median age of the subjects was 35 (Figure 2c).

### General psychophysics procedures

The subjects were tested in the Rockefeller University Hospital Outpatient Clinic. Psychophysical tests were self-administered and computerized using custom-written software applications that ran on netbooks. To prevent errors, all odor vials used in this study were barcoded. Subjects scanned each odor vial containing the stimulus before opening the vial, and were only prompted to proceed if the correct vial was scanned.

Subjects opened the vial, sniffed the contents, and were asked if they could smell anything. If the answer was “no,” they were directed to move on to the next stimulus. If the answer was “yes,” they were asked if they know what the smell is. If they answered “yes”, they were given a chance to describe the smell (Figure 8). Then they were asked a series of 23 questions about the smell (Figure 2a). For each question, they were presented with a slider that could be moved along a line. The final position of the slider was then translated into a scale from 0 to 100. The first three questions asked how familiar, strong, and pleasant the smell was. For these three questions, the slider started in the middle of the line (position 50) and subjects were required to move it. The other 20 questions were how well each of 20 descriptors (“edible”, “bakery”, “sweet”, “fruit”, “fish”, “garlic”, “spices”, “cold”, “sour”, “burnt”, “acid”, “warm”, “musky”, “sweaty”, “ammonia/urinous”, “decayed”, “wood”, “grass”, “flower”, “chemical”) applied to the smell. For these questions, the slider started at the bottom of the line (position 0) and subjects were not required to move it. The 20 descriptors were chosen because they are broad enough to be applied to enough stimuli in our set to allow for the development of models that predict the application of the descriptor based on molecular features. Other descriptors such as “pineapple” “cork” “wet paper” are so specific that they are applied to relatively few molecules [20].

Each subject came to the Rockefeller University Outpatient Clinic for 10 visits. During each visit, 100 stimuli were profiled. The order of stimuli was randomized differently for each subject. Although there are likely sequence effects for individual ratings (for example, a moderately pleasant odor is probably rated as more pleasant when it follows a series of very unpleasant odors than when it follows a series of several very pleasant odors), these are averaged out in the pooled data. Subjects carried out the study at their own pace with the typical pace of 1 stimulus/minute.

### Stimuli

Stimuli were presented in vials as 1 mL of the diluted molecule in paraffin oil. Information about the stimuli and their dilutions can be found in Additional File 1. The chemicals were >97% pure with a median purity of 98%. This is a limitation of this dataset because 3% impurity can have an impact on the percept, especially when the molecule itself is odorless, but the impurity has a smell.

### Molecular features

Molecular complexity (Figure 1b and Figure 9c, top) was computed using the Bertz/Hendrickson/Ihlenfeldt formula [23]. It is a rough estimate of the complexity of a molecule, and considers the variety of elements in the molecule as well as structural features including symmetry. Stereochemistry is not used as a criterion. In general, large compounds are more complex than small compounds. The correlation between molecular complexity and number of atoms (excluding H) among the 480 molecules used here is 0.88, but high symmetry and the lack of diversity in atom types results in lower complexity. The complexity values were obtained from PubChem. Vapor pressures (Figure 9b, top) were assembled from a variety of online sources, and were either experimentally measured or calculated.

Molecular features (Figure 9a) were calculated using Dragon 6 software (Talete) [26, 27]. Of the 4,885 molecular features, only the following categories were included in the analysis presented here: atom-centered fragments (115 descriptors), constitutional indices (43 descriptors), functional group counts (154 functional descriptors), molecular properties (20 descriptors), and ring descriptors (32 descriptors). Topological indices, walk and path counts, connectivity indices, information indices, 2D matrix-based descriptors, 2D autocorrelations, Burden eigenvalues, P_VSA-like descriptors, ETA indices, edge adjacency indices, geometrical descriptors, 3D matrix-based descriptors, 3D autocorrelations, RDF descriptors, 3D-MoRSE descriptors, WHIM descriptors, GETAWAY descriptors, Randic molecular profiles, atom-type E-state indices, CATS 2D, 2D atom Pairs, 3D atom Pairs, charge descriptors, and drug-like indices were not included in the analysis. Molecular features that had the same value for more than 98% of the molecules used here were also excluded from the analysis.

### Word cloud

The word cloud in Figure 8a shows how frequently certain words were used by the subjects to describe the smells of the stimuli. It was produced with the Wordle program at http://www.wordle.net and represents the frequency of word usage by font size. The program was set to remove common English words (“and” “but” “or” etc.), and the following words were manually excluded because they did not describe perceptual qualities: “smell” “smells” “smelly” “smelling” “odor” “sort” “also” “something” “kind” “mixed” “maybe” “flavor” “flavored” “strong” “slightly” “mildly” “type” “background” “like” “used” “hint” “mild” “bit” “reminds” “mix” “scented” “faintly” “scent”.

### Supporting data

All raw are included within the article and in Additional file 1. This spreadsheet list the odors used in this study (source, name, CID, C.A.S. number; odor dilutions), subject demographic information (gender, race, ethnicity, age), and the psychophysical dataset used for all the analysis in this paper.

## Author contributions

AK carried out all the experiments and analyzed the data. AK and LBV together designed the experiments, interpreted the results, produced the figures, and wrote the paper. Both authors read and approved the final manuscript.

## Acknowledgements

We thank our research volunteers for their patience in smelling the 1,000 stimuli used in this study and Peggy Hempstead for overseeing data collection. We further thank Joel Mainland, Pablo Meyer, Kevin Lee, and members of the Vosshall laboratory for discussion and helpful comments on the manuscript. The staff of The Rockefeller University Hospital Outpatient Clinic provided invaluable support in performing the experiments. Chris Vancil provided custom programming for the Rockefeller University Smell Study smell testing computer interface. This research was supported in part by grant # UL1RR024143 from the National Center for Research Resources, National Institutes of Health. AK was supported by a Branco Weiss Science in Society Fellowship. LBV is an investigator of the Howard Hughes Medical Institute. The funders had no role in the design, collection, analysis, or interpretation of data.

